# Delta phase-dependent modulation of temporal predictions by parietal transcranial alternating current stimulation

**DOI:** 10.1101/2024.11.18.624126

**Authors:** Rebecca Burke, Alexander Maÿe, Jonas Misselhorn, Marina Fiene, Felix J. Engelhardt, Till R. Schneider, Andreas K. Engel

## Abstract

**Background:** Previous research has shown that temporal prediction processes are associated with phase resets of low-frequency delta oscillations in a network of parietal, sensory and frontal areas during non-rhythmic sensory stimulation. Transcranial alternating current stimulation (tACS) modulates perceptually relevant brain oscillations in a frequency and phase-specific manner, allowing the assessment of their functional qualities in certain cognitive functions like temporal prediction.

**Objective:** We addressed the relation between oscillatory activity and temporal prediction by using tACS to manipulate brain activity in a sinusoidal manner. This enables the investigation of the relevance of low-frequency oscillations’ phase for temporal prediction.

**Methods:** Delta tACS was applied over the left and right parietal cortex in two separate unimodal and crossmodal experiments. Participants judged either the visual or the tactile reappearance of a uniformly moving visual stimulus, which shortly disappeared behind and occluder. Using an intermittent electrical stimulation protocol, tACS was applied with six different phase shifts relative to sensory stimulation in both experiments while participants performed a temporal prediction task. Additionally, a computational model was developed and analysed to elucidate oscillation-based functional principles for the generation of temporal predictions.

**Results:** Only in the unimodal experiment, the application of delta tACS resulted in a phase-dependent modulation of temporal prediction performance. By considering the effect of sustained tACS in the computational model, we demonstrate that the entrained dynamics can phase-specifically modulate temporal prediction accuracy.

**Conclusion:** Our findings suggest that phase-specific neuromodulation through delta tACS can influence temporal prediction accuracy in a unimodal context. This provides support to the notion that low-frequency delta oscillations are of causal relevance for temporal prediction.

## Introduction

Temporal prediction involves the brain’s ability to anticipate the timing of future events. This ability contributes to numerous aspects of human behaviour, from simple sensory processing to complex decision-making. Neural underpinnings related to temporal prediction emphasise the role of phase alignment or entrainment of neural oscillations, particularly in the low-frequency delta band, as a key mechanism [Lakatos et al., 2008, Schroeder and Lakatos, 2009]. Studies have shown that delta oscillations align with the temporal structure of the incoming sensory information stream and that oscillatory phase plays a crucial role in determining the accuracy of responses [Arnal et al., 2015, Stefanics et al., 2010]. When events are highly temporally predictable, such as rhythmic patterns, the high-excitability phase of ongoing oscillations can be adjusted to align with the predicted onset of incoming information, enhancing neural excitability and thereby optimising behaviour [Gulberti et al., 2015, Kayser et al., 2010, Lakatos et al., 2008]. This phase alignment is strengthened when top-down resources are engaged, e.g., when expectation cues indicate the occurrence of relevant stimuli [Herbst et al., 2022, Stefanics et al., 2010]. Furthermore, oscillatory phase in one modality can be modified not only by events within the same modality, but also by events from different modalities [Bauer et al., 2018, Diederich and Colonius, 2012, Fiebelkorn et al., 2011, Lakatos et al., 2008, Romei et al., 2012]. Supporting this mechanism, we previously found a strong correlation between delta inter-trial phase consistency (ITPC) in the parietal cortices and performance in a unimodal and crossmodal temporal prediction task [Daume et al., 2021]. This study suggests that phase-alignment of endogenous oscillations underlies temporal prediction processes. Yet, causal evidence for the functional role of low-frequency phase in encoding the onset of relevant stimuli is missing. Here, we set out to investigate the causal role of oscillatory phase in temporal prediction. To achieve this, precise control over both oscillatory phase and sensory stimulation is required [de Graaf and Sack, 2014, Fiene et al., 2022, 2020, Sack, 2006]. Extending our previous work [Daume et al., 2021], we delivered sensory stimuli with precise timing during transcranial alternating current stimulation (tACS) with systematic phase offset. If temporal prediction performance was related to delta phase, we expected to see a fluctuation in performance depending on the tACS phase, leading to either enhancement or deterioration of performance (Figure 1). Our results from behavioural data and computational modelling show that unimodal temporal prediction is dependent on delta oscillatory phase.

**Figure 1:**
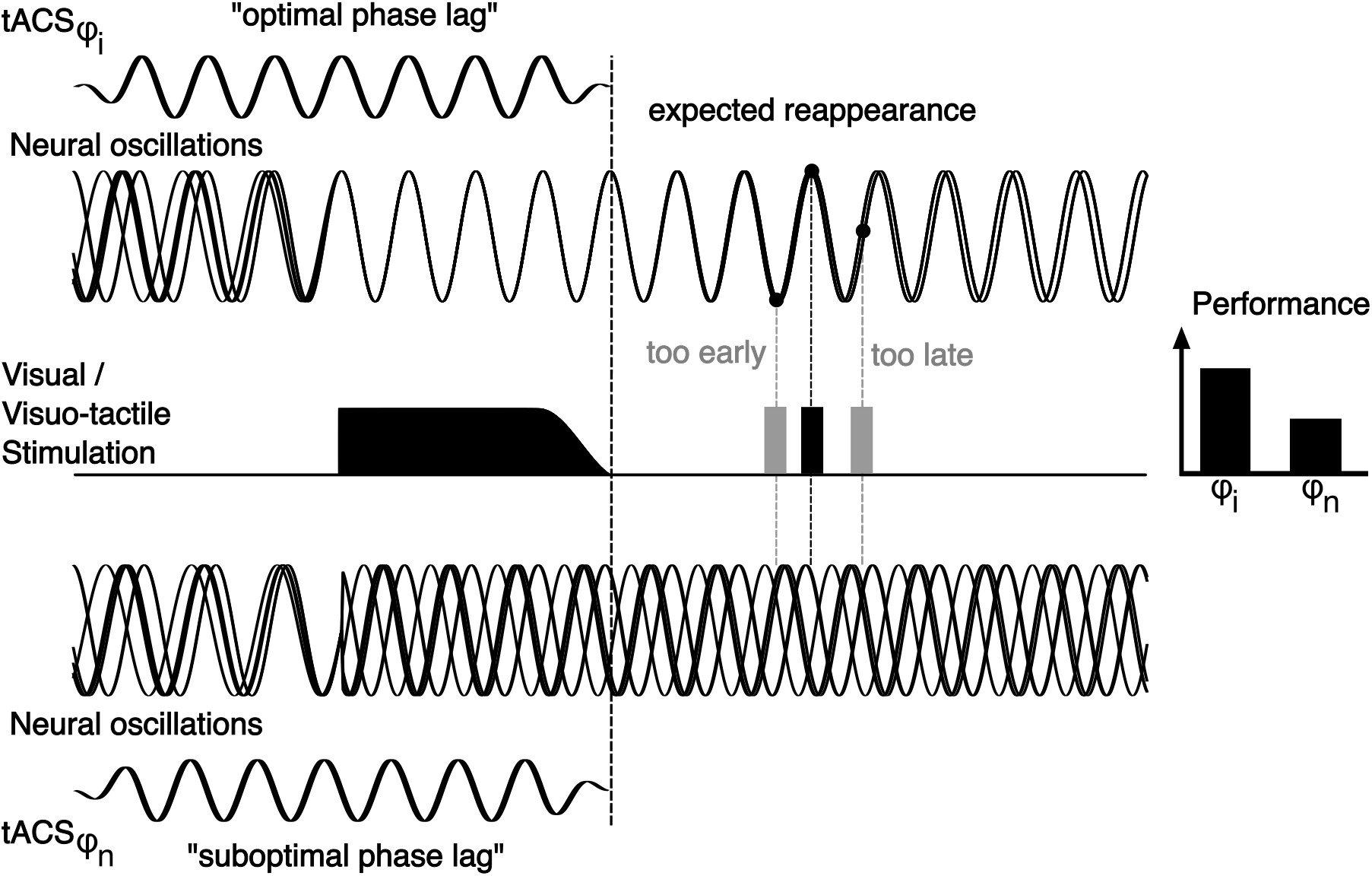
Rationale of the study. We explored how the timing of oscillatory phase affects temporal prediction by integrating sensory stimulation with transcranial alternating current stimulation (tACS). Building on Daume et al. [2021], our approach differed by using a gradually disappearing visual stimulus to reduce prominent sensory induced offset effects. Instead, the application of tACS was employed to induce phase alignment by engaging neural oscillations to synchronise with the electrical rhythm applied to the scalp. The notion of phase alignment assumes that the accuracy of predictions regarding incoming relevant information depends on the magnitude of phase alignment and that the phase encodes the expected time point of reappearance. The visual stimulus is designed to coincide with or disrupt the oscillatory phase alignment induced by tACS, thereby either maintaining the phase alignment or interfering with the ongoing oscillation. If the phase of neural oscillations constitutes an integral element for temporal prediction, we should either see an enhancement or a deterioration of performance depending on the tACS phase lag relative to the visual stimulus. Furthermore, an optimal phase lag of tACS would align the expected reappearance of the stimulus with a high-excitability phase of the neural oscillations and thereby augment temporal prediction performance.

## Material and methods

### Participants

In the unimodal experiment, 29 naïve, right-handed participants (mean age 26.5± 4.35 years, 16 female; 13 male) were recruited and in the crossmodal experiment, 30 naïve, right-handed participants (mean age 25.7 ±3.9 years, 15 female; 15 male) were recruited based on previous effect size observations in studies that investigated phase-dependent stimulation effects on sensory perception using tACS [Fiene et al., 2022, 2020]. Participants had normal vision and no psychiatric or neurological history. Handedness was assessed using the short version of the Edinburgh Handedness Inventory. The experiments, approved by the Hamburg Medical Association, adhered to the Declaration of Helsinki, with informed consent given by all participants and monetary compensation provided.

### Experimental procedure

The experimental design of the current studies was based on Roth et al. [2013] which examined visual temporal predictions in cerebellar patients, and closely aligns with a previous MEG study that we have carried out on the neural mechanisms of temporal predictions [Daume et al., 2021].

Participants sat in a dimly lit, electrically shielded and sound-attenuated chamber. The visual stimuli were projected onto a matte LCD screen at 120 Hz with a resolution of 1920 × 1080 px.

A white noise occluder with a red fixation dot (visual angle: 3.5° x 4.7° (h x w), corresponding to 150 × 200 px) processed with a Gaussian filter (imgaussfilt.m in MATLAB) was presented centrally against a grey background (Figure 2). Trials began with a fixation interval of 2000 ms accompanied by the application of tACS or active control stimulation, followed by a white ellipse stimulus (size: 1.7° x 0.5° visual angle) moving at 5.6°/s from the left periphery towards the occluder, and fading out over 500 ms before disappearing. After a certain time *t*, the visual stimulus reappeared on the right side of the occluder and continued moving for another 500 ms at the same constant velocity as before the occlusion (Figure 2A). In each trial, the reappearance time of the stimulus varied randomly between Δ*t* = ±17 and ±467 ms (±2 to ±60 frames; refresh rate:120 Hz) in steps of 50 ms (3 frames) relative to the objectively correct reappearance time of 1500 ms. Participants were instructed to judge whether the stimulus reappeared *too early* or *too late* via button press with their right hand (BlackBoxToolKit USB Response Pad, Black Box ToolKit Ltd) based on the stimulus’ movement before the occluder and the estimated movement behind the occluder. Response mapping of the two buttons was counterbalanced across all participants.

**Figure 2:**
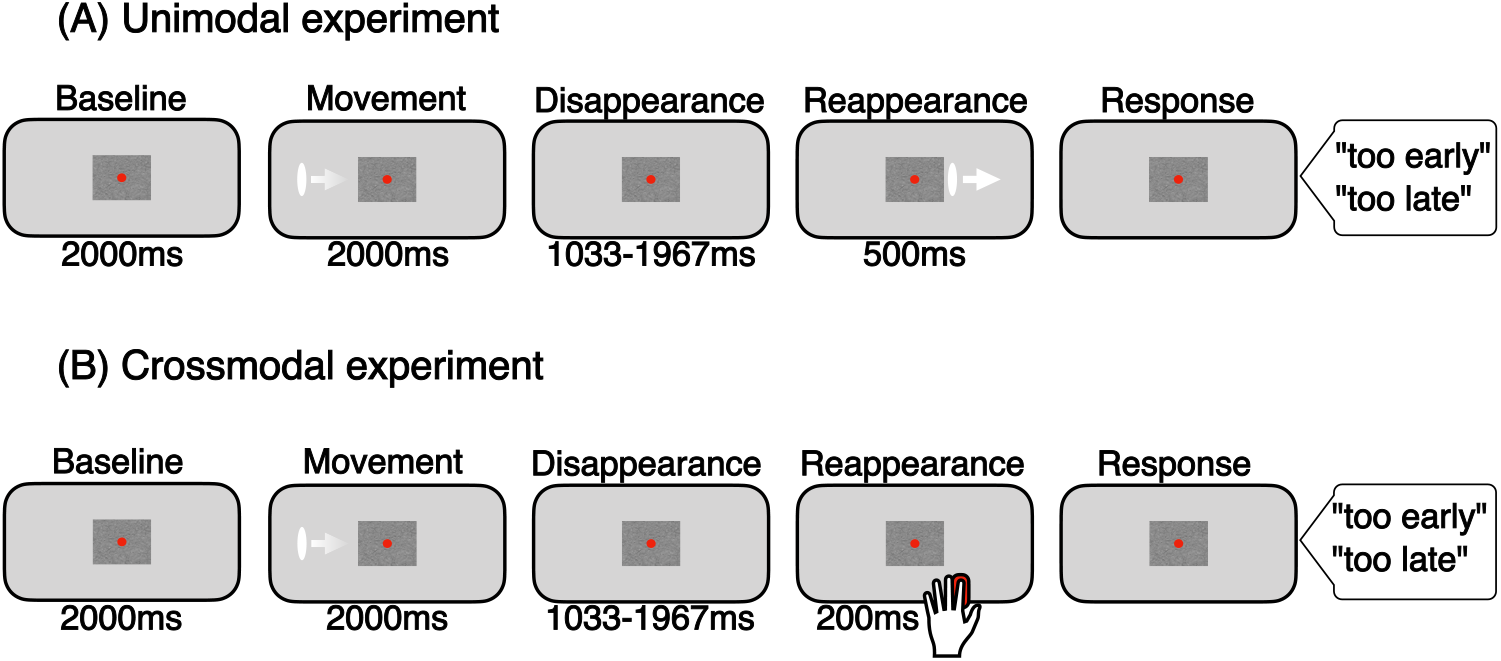
Experimental design. (A) Unimodal experiment: In the beginning of each trial, a white ellipse appeared on the screen and moves towards the occluder before disappearing. After 1500 ms *±*17 to *±*467 ms the stimulus reappears. The participants then had to decide whether the stimulus reappeared “too early” or “too late”. (B) Crossmodal experiment: Like in the unimodal experiment, the participants saw a white ellipse moving towards the occluder before disappearing behind it. After time intervals of 1500 ms *±*17 to *±*467 ms, the participants received a tactile stimulus, signalling the reappearance. Again, the participant’s task was to identify whether the tactile stimulus reappeared “too early” or “too late”.

The crossmodal experiment mirrored the unimodal setup, substituting the visual reappearance with a tactile stimulus to the left index finger (Figure 2B). The tactile stimulus indicated the time point *t±* Δ*t* (as described in the unimodal design) at which the disappearing visual stimulus would have reappeared on the other side of the occluder. The tactile stimulus was presented using a Braille piezo-stimulator (QuaeroSys, Stuttgart, Germany; 2 × 4 pins, each 1 mm in diameter with a spacing of 2.5 mm) by pushing all eight pins up simultaneously for 200 ms. Participants were instructed to respond as fast and as accurate as possible, once the visual or tactile stimulus reappeared after the occluder. Immediate feedback was not provided, but participants received block-wise performance summaries and could rest between blocks. Each participant took part in two sessions on two days, maintaining consistent timing of the day and response mapping. Each session comprised ten blocks of 60 trials, resulting in a total of 600 trials. Prior to the main experiment, participants carried out a training block of 30 trials to get familiar with the task and stimulus material and received immediate feedback about the correctness of their response. To mask the sound of the Braille stimulator during tactile stimulation and other background noise, participants wore ear plugs for the whole duration of the recording session in both experimental designs. At the end of the second recording day, participants filled out a questionnaire asking for a specific strategy they might have used for the temporal prediction task. We used MATLAB R2016b (MathWorks, Natick, USA; RRID: SCR_001622) and Psychophysics Toolbox (RRID: SCR_002881) [Brainard, 1997] on an MS-Windows 10 operating system for stimulus presentation.

### Transcranial alternating current stimulation

Participants attended two stimulation sessions in which two high-definition-tACS (HD-tACS) 2×2-montage (Figure 3B) were employed at a frequency of 2 Hz and a peak-to-peak current of

**Figure 3:**
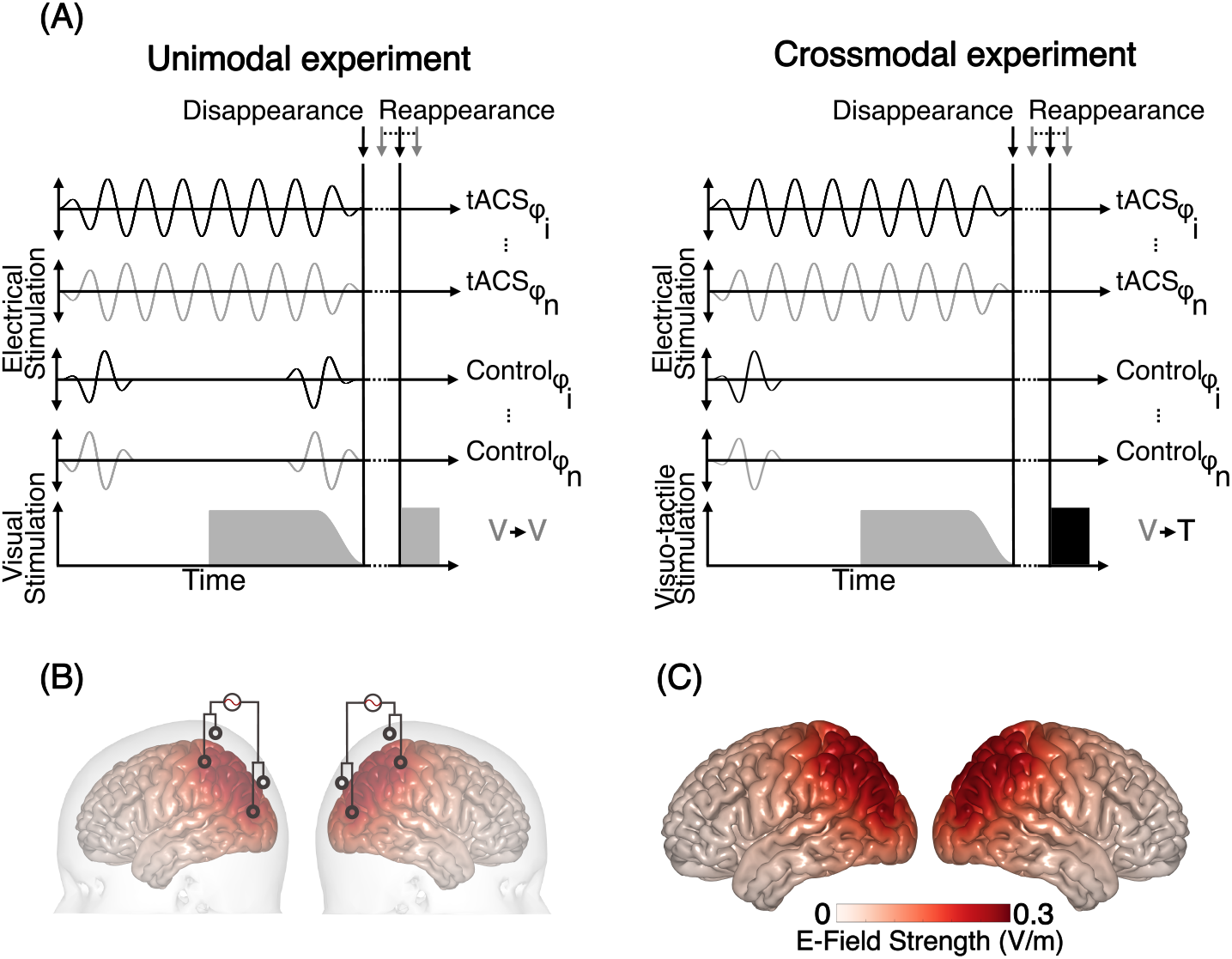
Experimental design and electrical field simulation. (A) tACS was applied at 2 Hz in a verum condition and an active control condition. In the verum condition, tACS was applied to the scalp intermittently for 4 s at six different phase lags (0°, 60°, 120°, 180°, 240°, 300°) relative to the visual stimulation. The active control condition of the unimodal experiment consisted of short ramps of tACS (1 s) in the beginning and in the end of the stimulation interval. In the crossmodal experiment, one second of ramped tACS was applied only in the beginning of the stimulation interval. In each trial of both experiments, a white ellipse was moving across the screen two seconds after tACS onset, then disappearing behind an occluder for various time intervals. The reappearing stimulus was either a visual or a tactile stimulus for the unimodal and crossmodal experiment, respectively (see Figure 2). (B) In-phase high-definition tACS montage targeting the left and right parietal cortices. (C) Simulated electrical field generated by the 2×2-montage with a peak-to-peak current of 2mA.

2 mA to target the left and right parietal cortex. The electric field 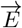 was estimated prior to the experiment by linear superposition of lead fields 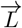,weighted by the injected currents *α*_*i*_ at stimulation electrodes *i* = {1,2,3,4}:

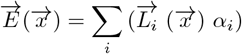

The lead field matrix 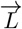 was computed by using boundary element three-shell head-models [Nolte and Dassios, 2005]. The estimated electrical field intensities were comparable to those shown to effectively modulate neural activity in both non-human primates and humans [Huang et al., 2018, Johnson et al., 2020, Kasten et al., 2019] (Figure 3C). Additionally, the tACS montage was slightly modified between the two experiments to account for differences between unimodal and crossmodal stimulation [Daume et al., 2021]. Despite this adjustment, the electrical fields remained comparable across both experiments.

The stimulation was administered using 12 mm Ag/AgCl electrodes and neuroConn stimulators (DC-Stimulator plus, neuroConn, Illmenau, Germany) with Signa electrolyte gel (Parker Laboratories Inc.) to maintain electrode impedances below 25 *k*Ω and ensure uniform electric fields. To minimize transcutaneous effects, EMLA cream (2.5% lidocaine, 2.5% prilocaine) was applied for local anaesthesia one hour prior to the experiment.

tACS was delivered intermittently, with 4 s intervals and current ramps of 500 ms at six different phase lags relative to the visual stimulation (0°, 60°, 120°, 180°, 240°, 300°). In the unimodal design’s active control condition, the current was ramped in and out over 1 s at both interval ends, whereas in the crossmodal design, the ramp occurred only at the interval start (Figure 3A). Post-session, a questionnaire assessed perceived intensity and temporal evolvement of side effects like skin sensations, fatigue, and phosphene perception.

To ensure participants did not use potential rhythmic tactile sensations resulting from tACS for temporal prediction, they completed a tapping test after the second session. Participants were exposed to 4s intervals of tACS, matching the tACS rhythm used in the experiment, and were instructed to tap in rhythm with the stimulation if possible. Afterwards, having been shown the 2 Hz rhythm by the experimenter tapping on a table, they repeated the test. This test aimed to determine whether participants could perceive the stimulation while actively attending to it, ruling out the possibility that positive tACS effects were solely due to transcutaneous/somatosensory effects rather than direct neural effects [Asamoah et al., 2019]. Two participants were able to tap in the presented rhythm and were therefore excluded from further analyses. In the unimodal experiment, 2 out of 29 participants tapped correctly to the 2 Hz rhythm initially and were removed from data analysis; in the crossmodal experiment, 4 out of 30 participants did. Notably, detailed instructions about the 2 Hz rhythm did not improve participants’ ability to tap in sync.

### Statistical analysis of behavioural data in relation to the tACS phase

Behavioural effects of tACS were evaluated by calculating accuracy values for the six different phase bins. The tACS-phase-specific performance modulation was quantified using two measures.

Firstly, the Kullback-Leibler divergence (DKL) was calculated, which quantifies the deviation of the observed amplitude distribution (P) from a uniform distribution (Q) defined as

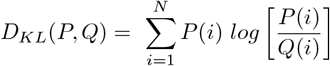

where N is the number of phase bins [Tort et al., 2010]. DKL values can range from 0 (indicating a strictly uniform distribution) to 1 (indicating a nonuniform distribution). Therefore, as the distance between P and Q increases, the DKL value also increases. The input to this function were the accuracy values normalised by the sum of the performance values across all six conditions.

Additionally, it was examined whether the performance modulation induced by tACS over phase conditions follows a cyclic pattern, as expected due to the sinusoidal nature of tACS. A one-cycle sine function was fit to the raw accuracy values of each participant using a nonlinear least-squares algorithm,

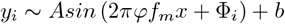

where *y*_*i*_ is the observed performance. *A*, Φ_*i*_, and *b* represent the amplitude, the phase lag of the sine fit, and the intercept, respectively. For each participant, the best sine fit function *g*_*i*_(*x*) = *Asin* (2*πf*_*m*_*x* + Φ_*i*_) + *b* was obtained for each condition (tACS and active control) separately by applying the nonlinear least-squares algorithm, allowing amplitude, phase lag and intercept to vary (three degrees of freedom). From this fit, the amplitude value was extracted. The goodness of fit (*R*^2^) of the resulting best fitting function was calculated for each participant. Therefore, while DKL provides a general measure of temporal prediction performance modulation, the sine fit allows an assessment of systematic modulation patterns that depend on the tACS phase.

Statistical significance of DKL and sine fit amplitude value differences between the verum and active control condition were examined using paired-samples t-tests. Effect sizes were calculated using Cohen’s *d*_*z*_.

Since traditional paired-samples t-tests are not sufficient to interpret null results, we used Bayesian hypothesis testing [de Graaf and Sack, 2018, Kass and Raftery, 1995]. This approach incorporates prior beliefs and the observed data to assess the evidence for or against the null hypothesis. This analysis produces a single value known as the Bayes Factor (*BF* _10_*/BF* _01_). In essence, it determines which hypothesis aligns better with the data and to what degree, indicating which hypothesis provides a more accurate prediction of the observed data. For this analysis the MATLAB package Bayes Factor was used [Krekelberg, 2022].

### Modelling temporal predictions

#### Oscillator ensemble model

To deepen understanding of neuronal mechanisms for temporal predictions, we developed a computational model inspired by previous lab work [Maye et al., 2019] but with a new mathematical approach focusing on temporal dynamics. The model uses Hopf oscillators, modified by Hebbian learning for frequency tuning [Righetti et al., 2006], to adapt to periodic inputs.

The Hopf oscillator is composed of an excitatory element *x* and an inhibitory element *y*:

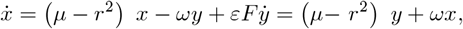

where *r*^2^ = *x*^2^ + *y*^2^, *ω* is the intrinsic oscillation frequency, and *µ* defines the radius of the limit cycle, i.e., the oscillator’s amplitude. The parameter *ε* controls how strongly an external input *F* affects the intrinsic dynamics of the oscillator. This oscillator can adjust its frequency to a periodic input signal by means of the following learning rule [Righetti et al., 2006]:

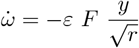

An ensemble of these adaptive-frequency oscillators (*N* =100) with random initial conditions learns to decompose a periodic input signal into its various components. Here, a collective behaviour was expected in which oscillators tune to one of the input components if the frequencies are close but continue their intrinsic dynamics if they are not.

The model can be linked to neurophysiological processes by considering *x* and *y* as a description of the various excitatory-inhibitory circuits that have been found in the visual cortex [Adini et al., 1997, Donner and Siegel, 2011, Xu et al., 2016]. The input *F* = sin *ω*_*sens*_ conceives a motion-induced oscillation that has been described in visual cortices of cats [Gray and Viana Di Prisco, 1997] and humans [Orekhova et al., 2015].

#### Modelling tACS

Our model assumes that tACS might modulate the coupling strength of the sensory input to the excitatory-inhibitory circuits. Formally this modulation can be modelled by a sinusoidal tACS signal and an additional coupling parameter *κ* which can be used to control the modulation depth:

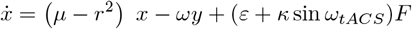

The parameters were set as follows: *µ* = 1, *ε* = 0.1, *κ* = 0.1, *ω*_*tACS*_ = 0.2*π*, and *ω*_*sens*_ = 4*π*. The initial conditions were randomized in the intervals *x, y* ∈ [−1, 1] and *ω* ∈ [2*π*, 6*π*]. The equations were numerically solved by a 4^th^ order Runge-Kutta method in the interval *t* ∈ [0, 100] with *dt* = 0.01 as the temporal resolution. The input *F* was switched off for 2s at the end of the stimulation period to reflect the occlusion of the stimulus during the prediction period.

Over time, oscillators adjust their frequencies to match sensory inputs. Research indicates stronger ITPC during the time window of occlusion [Daume et al., 2021]. In line with the idea that oscillatory phase encodes the time point of reappearance through alignment of high excitability phases to the predicted onset of reappearance, this phase consistency should persist at object reappearance. To test this hypothesis, the model sampled phase distributions at the stimulus’ reappearance through multiple trials with randomised initial conditions during a training phase of the model. The average of the phases *Φ*_*n*_ across the oscillator ensemble was considered as the ensemble’s phase Φ:

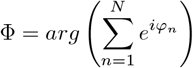

This distribution corresponds to subjectively correct reappearance of the stimulus. In the test phase, ensemble phases tied to the time windows corresponding to earlier or late reappearances were sampled. These phases were then compared against the learned phase distribution for right-on-time reappearances. If most phases exceeded the test phase, the model indicated that the stimulus reappeared *too late* and vice versa.

## Results

### Phase-specific modulation of unimodal temporal prediction

Averaged across phase lags, participants executed the temporal prediction task better than chance, but below ceiling 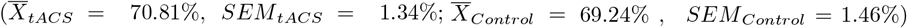. To assess the phasic modulation of temporal prediction performance by tACS, we quantified the behavioural modulation strength by two measures. First, the DKL measure shows that the behavioural performance across phase lags is nonuniformly distributed. The strength of this modulation was significantly greater during tACS compared to the active control condition (*t*(27) = 2.829, *p* = .008, *d*_*z*_ = 0.54) (Figure 4A).

**Figure 4:**
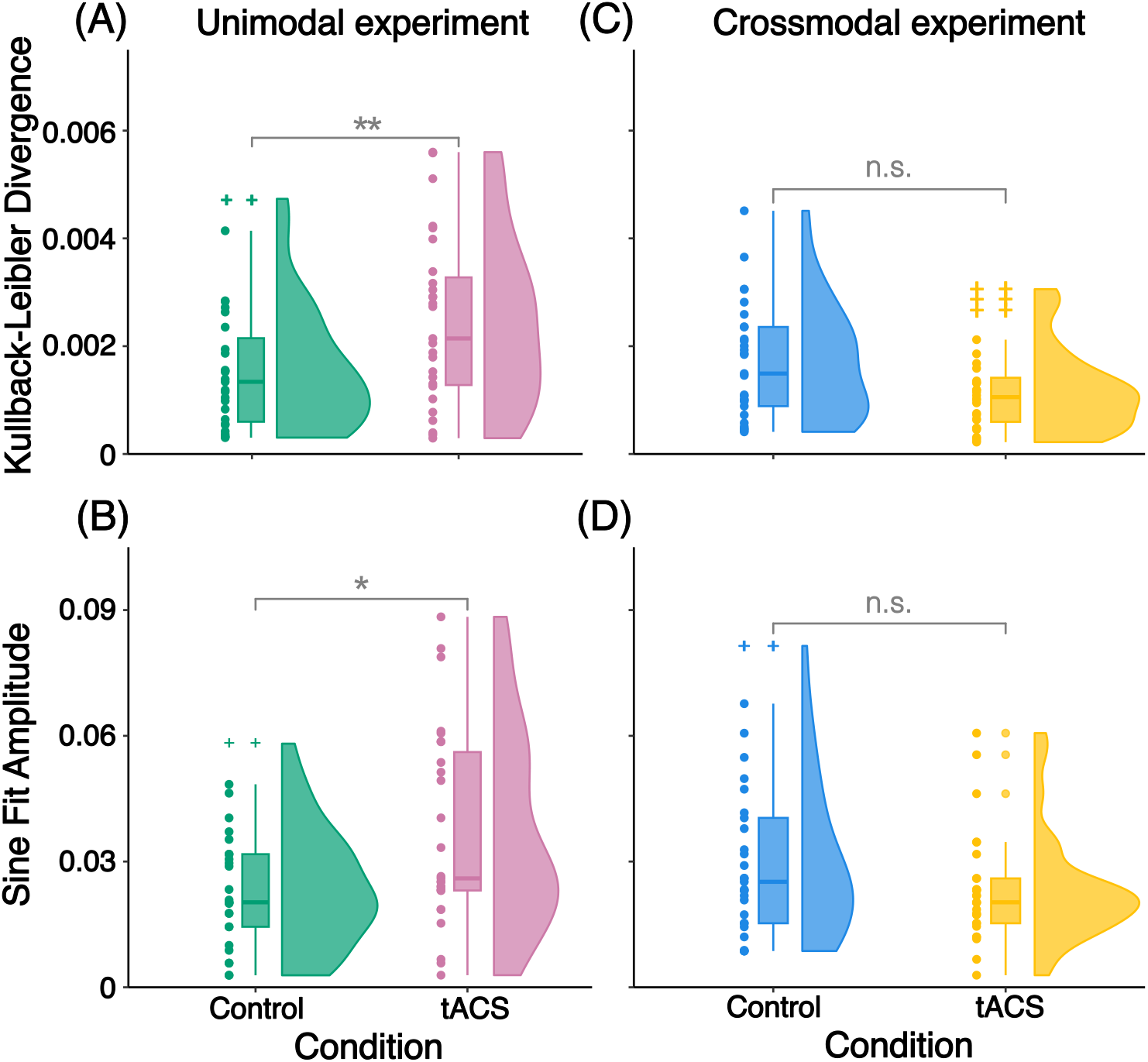
Temporal prediction performance modulation quantified via two modulation measures. (A, B) Significantly increased Kullback-Leibler divergence (DKL) and sine fit amplitudes under tACS relative to active control condition in the unimodal experiment. (C, D) No significant difference in DKL values and sine fit amplitudes under tACS compared to the active control condition in the crossmodal experiment. Asterisks mark the uncorrected p-value of dependent samples t-tests; *p < .05, **p<.01.

Second, to check whether behavioural performance systematically fluctuated around the mean along the 2 Hz tACS phase, we fitted sinusoidal waves to the behavioural performance across tACS phase lags relative to the offset of the visual stimulus. Figure 5A shows the behavioural data of all 27 participants for both, verum and active control tACS. Comparing the amplitude values obtained from the most suitable sinusoidal curve for both conditions revealed a significantly higher modulation strength in the tACS condition compared to the active control condition (one-sided paired-samples t-test: *t*(27) = 2.543, *p* = .01, *d*_*z*_ = 0.49) (Figure 4B).

**Figure 5:**
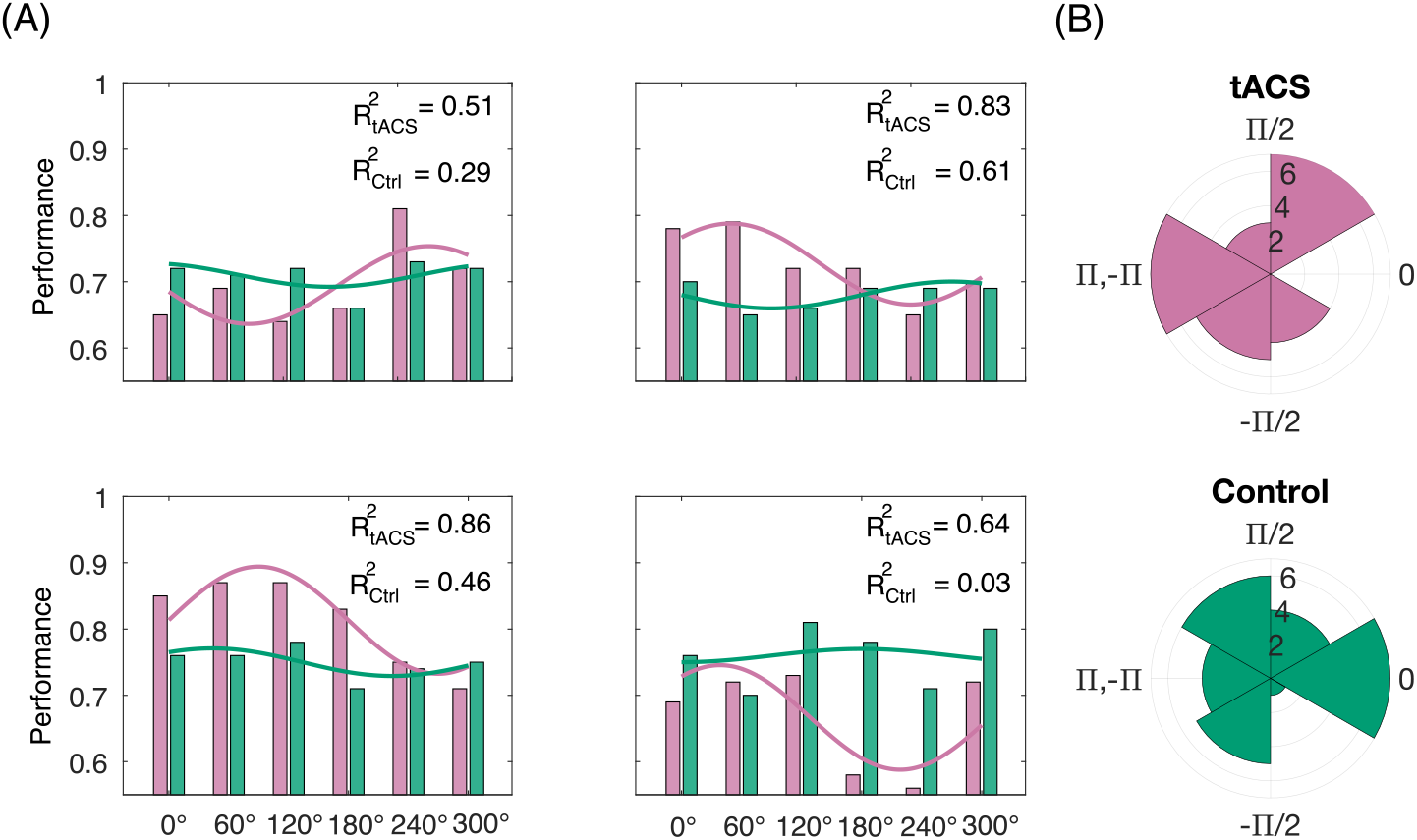
(A) Sine fits of 4 exemplary subjects for unimodal temporal prediction experiment showing the temporal prediction performance for the six phase-lags (green: Control, red: tACS) and (B) the distribution of the individual optimal phase lags of tACS and control condition in the unimodal experiment.

Although the observed relation between tACS phase and behavioural performance was robust, there was significant variability in the optimal phase among participants. Thus, different participants reached their peak behavioural performance at different tACS phase lags. In fact, when considering the entire sample (N=27), there was no evidence of a distinct concentration of behavioural entrainment at a specific phase (Rayleigh test for nonuniform distribution: *z*_*tACS*_ = 1.04, *p*_*tACS*_ = 0.36; *z*_*Control*_ = 0.44, *p*_*Control*_ = 0.65, see Figure 5B).

Together, we observed a significant impact of tACS phase on temporal prediction performance in the unimodal temporal prediction task. These results suggest a substantial influence of neural low-frequency oscillatory phase on unimodal temporal prediction.

### tACS does not phase-specifically modulate crossmodal temporal prediction

For the crossmodal temporal prediction paradigm, we could not find a similar relationship between the tACS and visuo-tactile stimulation. When statistically testing for differences between the two tACS conditions, we could neither find a significant difference in the analysis of general phasic modulation (*DKL* : *t*(26) = −0.807, *p* = .427, *d*_*z*_ = −0.149), nor could we find a significant effect in the analysis of sinusoidal modulation (*Sine F it* : *t*(26) = −1.478, *p* = .151, *d*_*z*_ = −0.274) (Figure 4C/D).

To assess the evidence for the null hypothesis and, therefore, allowing an interpretation of the null results, the Bayes Factor was calculated (*DKL* : *BF* _01_(26) = 3.76). This value indicates that the current data is 3.76 times more likely to be observed if the null hypothesis was true (i.e., no general phasic modulation through tACS compared to the active control condition), than if the alternative hypothesis is true. According to recommended guidelines, this provides “moderate” evidence in support of the null hypothesis (BF>3, but <10). Furthermore, in the analysis of sinusoidal modulation of temporal prediction performance (*Sine F it* : *BF* _01_ = 1.91), the Bayes Factor can be descriptively interpreted as providing merely anecdotal evidence for the null hypothesis to be true (BF<3).

### tACS-entrainment in the computational model

Under the influence of the motion-induced oscillatory input to the model, the initially random phase distribution of the ensemble developed towards a predominant phase of approximately *π* at the time point of the actual reappearance of the moving object (Figure 6A, top). This shift towards a predominant phase corresponds to the subject-specific optimal phase in the behavioural data of our experiment. In addition, oscillators with initial frequencies close to the input frequency also adjusted their frequencies to the input (Figure 6A, bottom).

**Figure 6:**
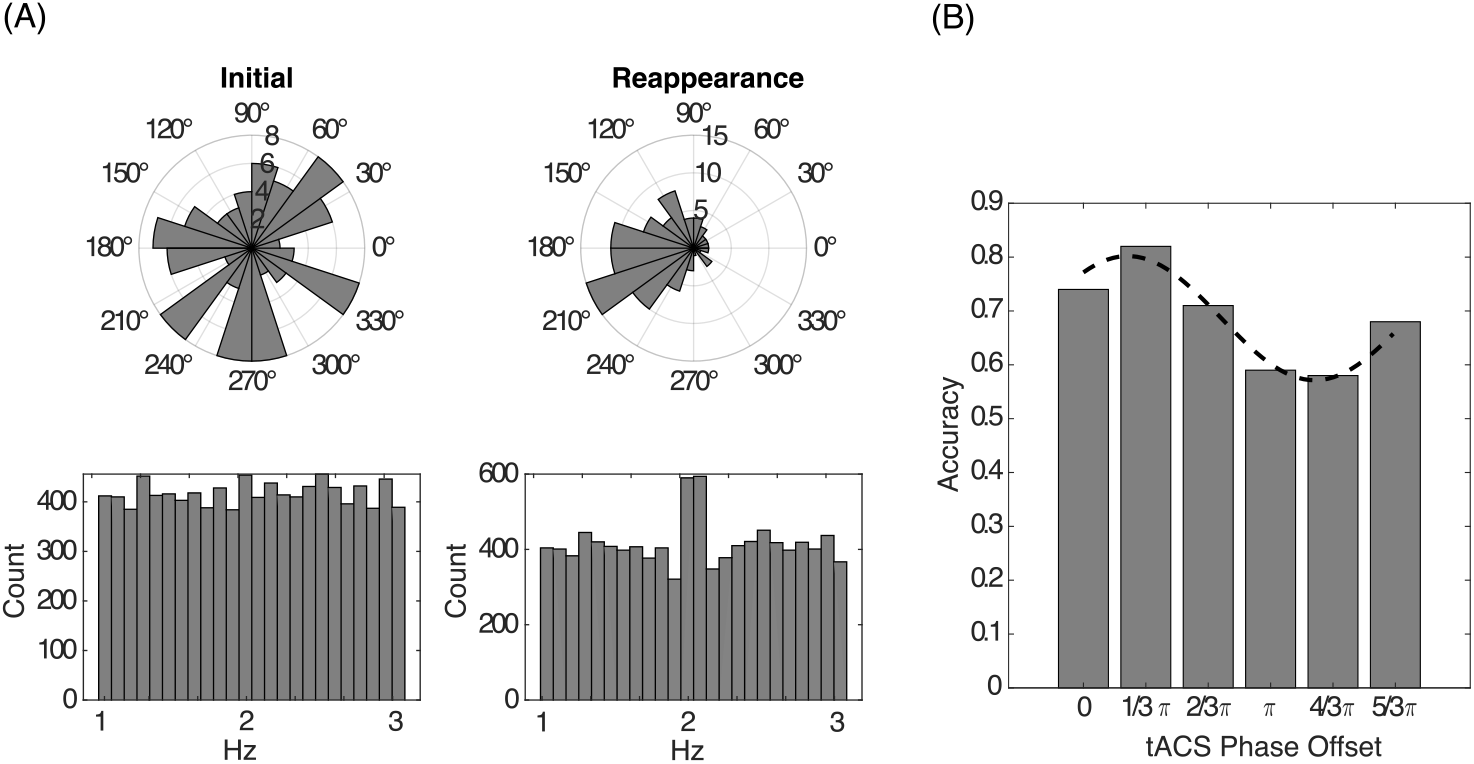
(A) Shows exemplary model data without tACS input. Histograms of phases (top) and distribution of intrinsic oscillation frequencies *ω* (bottom) of the ensemble oscillators at t=0 (left) and at the reappearance (right) of the moving object. Data are from 100 repetitions of an ensemble of 100 oscillators. (B) Exemplary temporal prediction accuracy of the model for different tACS phase offsets. The dashed line shows a fitted sine wave.

The parameter *ε* was adjusted to match the overall accuracy of the model’s temporal prediction accuracy approximately to the results from the behavioral study. The simulation of different tACS phase offsets relative to the motion-induced input resulted in the same characteristic sinusoidal modulation of temporal prediction accuracies like in the participants (Figure 6B).

## Discussion

The present studies aimed to investigate the causal role of oscillatory phase in temporal prediction. We employed an intermittent electrical stimulation design in which we systematically combined rhythmic electrical with continuous visual or visuo-tactile stimulation. We hypothesized that applying tACS at six different phase lags relative to the disappearance of a continuously moving visual stimulus would result in a phase-dependent modulation of temporal prediction of the visual stimulus’ reappearance. Our results confirm that unimodal visual temporal prediction performance was modulated in a tACS phase-dependent manner, leading to both enhancement and deterioration of performance. Under the assumption that neural oscillations were entrained by tACS, our data suggests a functional role of parietal low-frequency neural oscillations for temporal prediction. The computational model showcases that neural oscillations influenced by visual motion causes the initially random phase distribution to converge toward a predominant phase, thereby essentially increasing ITPC. Additionally, incorporating phase-specific tACS parameters into the model allowed us to show that motion-induced neural oscillations, in relation to the tACS phase, either preserves or disrupts the phase alignment induced by tACS and thereby effectively enhances or deteriorates the temporal prediction accuracy of the model. Our model suggests a stronger coupling between the sensory input and the oscillator dynamics as a potential entrainment mechanism.

### Implications for understanding the neural mechanisms of temporal prediction

tACS offers a distinctive approach to explore and confirm hypotheses on targeted brain regions, frequency bands and oscillatory phase. Although the sources of oscillatory activity may vary by temporal prediction task, the parietal cortex has been implicated in regulating temporal prediction across various tasks [Adini et al., 1997, Bueti et al., 2010, Bueti and Walsh, 2009, Cotti et al., 2011, Daume et al., 2021, Davranche et al., 2011, Gontier et al., 2013, Lewis and Miall, 2006, Mioni et al., 2020, Rao et al., 2001]. Hence, the parietal cortex has been a popular target for non-invasive brain stimulation when investigating temporal processing [Mioni et al., 2020]. In Alexander et al. [2005], the application of low frequency (1 Hz) repetitive transcranial magnetic stimulation (rTMS) to the parietal cortex led to a deterioration of performance in a temporal judgement task. Studies have revealed a hemispheric lateralisation for temporal versus spatial orienting in left and right parietal cortices, respectively [Coull and Nobre, 1998]. Consistent with these findings, Javadi et al. [2014] observed that left cathodal tDCS of the parietal cortex improved performance in duration judgment, aligning with findings by Rao et al. [2001] that the parietal cortex is crucial for encoding or maintaining temporal intervals. Here, we were able to demonstrate the significance of parietal low-frequency oscillatory phase for visual temporal prediction.

Previous research has highlighted the significance of neural oscillations in the delta range (0.5-3 Hz) for temporal prediction [Arnal and Giraud, 2012, Ng et al., 2013]. Entrainment through alignment of low-frequency delta oscillations with external cues is acknowledged as an important and flexible mechanism for improving sensory processing [Besle et al., 2011, Cravo et al., 2013, Lakatos et al., 2008, 2009]. Moreover, it has been suggested that the functional role of low-frequency delta oscillations in human anticipatory mechanisms involves the modulation of synchronized rhythmic fluctuations in the excitability of neuronal populations. When additional top-down resources are employed, this synchronisation through phase reset is enhanced [Herbst et al., 2022, Stefanics et al., 2010]. The phase of low-frequency delta oscillations has further been linked to accuracy in temporal judgment tasks [Arnal et al., 2015] and to precise behavioural adjustments made subsequent to temporal errors [Barne et al., 2017]. Here, we extend the findings of Daume et al. [2021] by causally inferring with tACS that the phase of aligned low frequency delta oscillations to the external stimulus encodes the onset of upcoming information through alignment of high-excitability phases of the oscillation with the time window of expected reappearance. This alignment allows optimal sampling of incoming information, considering top-down expectations regarding the occurrence and timing of stimuli [ten Oever et al., 2015].

Notably, the temporal structure (i.e., the velocity of the moving stimulus and the time window of disappearance) in our experiments has been restricted to the delta frequency range. This aligns with the frequency range of endogenous brain oscillations expected to be functionally relevant for temporal prediction [Lakatos et al., 2008]. It remains to be explored whether the observed sinusoidal modulations of behavioural performance in the unimodal task can occur outside of this frequency range. This could be investigated with different velocities of the fading visual stimulus and different temporal scales of the disappearance window to gain a comprehensive understanding of how oscillatory frequency and phase influences temporal prediction.

Each participant reached their peak performance at different phase lags, suggesting individual differences in the effects of tACS. The heterogeneity could be attributed to varying levels of neuronal entrainment to the 2 Hz tACS rhythm due to interindividual differences in intrinsic delta rhythms as well as to variability in anatomical connectivity and cortical morphology, leading to different neural phase lags [Bauer et al., 2018, Besle et al., 2011, Henry and Obleser, 2012, Kösem et al., 2014]. Furthermore, the variable effects of tACS may be due to different field orientation and variability in electrical current intensity concerning the target area. A prospective approach could involve not only considering individual stimulation location but also optimising individual current intensities to optimise tACS efficacy [Radecke et al., 2020]. Finally, the application of tACS at a fixed frequency suggests that entrainment effects might be comparatively weaker and tailoring the stimulation to the individual frequency may enhance tACS effects [Lorenz et al., 2019].

### Comparison of unimodal visual and crossmodal visuo-tactile temporal prediction

The crossmodal experiment revealed no significant impact of tACS on temporal prediction. Our Bayesian analysis provided weak to moderate support for this null effect, suggesting that our non-significant findings, in fact, might be merely coincidental [Ronconi et al., 2022]. Crossmodal sensory integration for shaping temporal predictions may have increased task difficulty, resulting in a greater variability in performance and thereby potentially diminishing small tACS effects. It was shown that assessing the duration of empty time intervals between different sensory modalities is harder than within the same modality, leading to decreased performance. This diminished performance was attributed to a suggested attentional bias induced by the cognitive load and the requirement to switch between modalities [Gontier et al., 2013]. Moreover, the electrical field of the tACS montage, although slightly adjusted, did not differ significantly between the crossmodal and unimodal experiment. However, the crossmodal condition is likely to involve at least partially different cortical areas and, thus, more specific optimization might be required to achieve phase-specific effects in the crossmodal condition. Furthermore, crossmodal temporal prediction might also involve frequency bands beyond the scope of our study. Several studies noted that building temporal predictions of upcoming events can be attributed to alpha-band desynchronisation and/or beta-band modulation [Arnal and Giraud, 2012, Daume et al., 2021, van Ede et al., 2011]. Desynchronisation of the respective frequency-band activity in these studies was suggested to serve as an active inhibitory mechanism that gates sensory information processing as a function of cognitive relevance. Therefore, the current study design cannot discern whether the noted disparities between the results of the unimodal and crossmodal experiment originate from variations in how oscillatory activity affects temporal prediction across different sensory modalities or are a result of discrepancies inherent in the two experimental designs. Future research on crossmodal entrainments may shed light on the underlying mechanisms by incorporating both unimodal and crossmodal temporal prediction tasks within a single study and by recording neurophysiological data to address the issue of neural efficacy [de Graaf and Sack, 2018].

## Conclusion

In the present experiments, we used tACS over bilateral parietal cortex to examine the causal relevance of delta oscillations for temporal prediction. In the unimodal task, we found a significant behavioural modulation of prediction accuracy in line with a phasic modulation of prediction-related neural activity. These behavioural results were replicated in a computational model which suggests that phase alignment sinusoidally fluctuates under the influence of phase-specific tACS. Accordingly, predictions about relevant future events can be implemented by phase adjustments to enhance sensory processing at the predicted time, leading to effective ‘gating’ of sensory inputs. The notion of a simple, generic mechanism of parietal delta phase for temporal prediction is challenged by our results from the crossmodal study in which phasic behavioural modulation by tACS was absent. We propose that future experiments should address this disparity in findings by exploring possible effects of task difficulty and oscillatory frequency on temporal prediction.

## CRediT authorship contribution statement

**Rebecca Burke:** Conceptualisation, Methodology, Investigation, Software, Formal analysis, Validation, Visualisation, Data curation, Writing - original draft, Writing - review & editing. **Alexander Maÿe:** Methodology – Computational Modelling, Writing – review & editing. **Jonas Misselhorn:** Conceptualisation, Methodology, Writing - review & editing. **Marina Fiene:** Conceptualisation, Methodology, Writing - review & editing. **Felix J. Engelhardt:** Conceptualisation, Investigation, Writing – review & editing. **Till R. Schneider:** Conceptualisation, Methodology, Writing - review & editing, Supervision. **Andreas K. Engel:** Conceptualisation, Methodology, Writing - review & editing, Funding acquisition, Project administration, Supervision.

## Acknowledgements

This work was supported by grants from the Deutsche Forschungsgemeinschaft (SFB TRR 169/B1/B4 awarded to A.K.E.) and from the European Union (project cICMs, ERC-2022-AdG-101097402 awarded to A.K.E.). We thank Karin Reimann for assistance in data recording, and Marleen J. Schoenfeld and Peng Wang for helpful discussions on the data. Views and opinions expressed in this paper are those of the authors only and do not necessarily reflect those of the European Union or the European Research Council. Neither the European Union nor the granting authority can be held responsible for them.

