## Supplementary Material for "Delta phase-dependent modulation of temporal predictions by parietal transcranial alternating current stimulation"

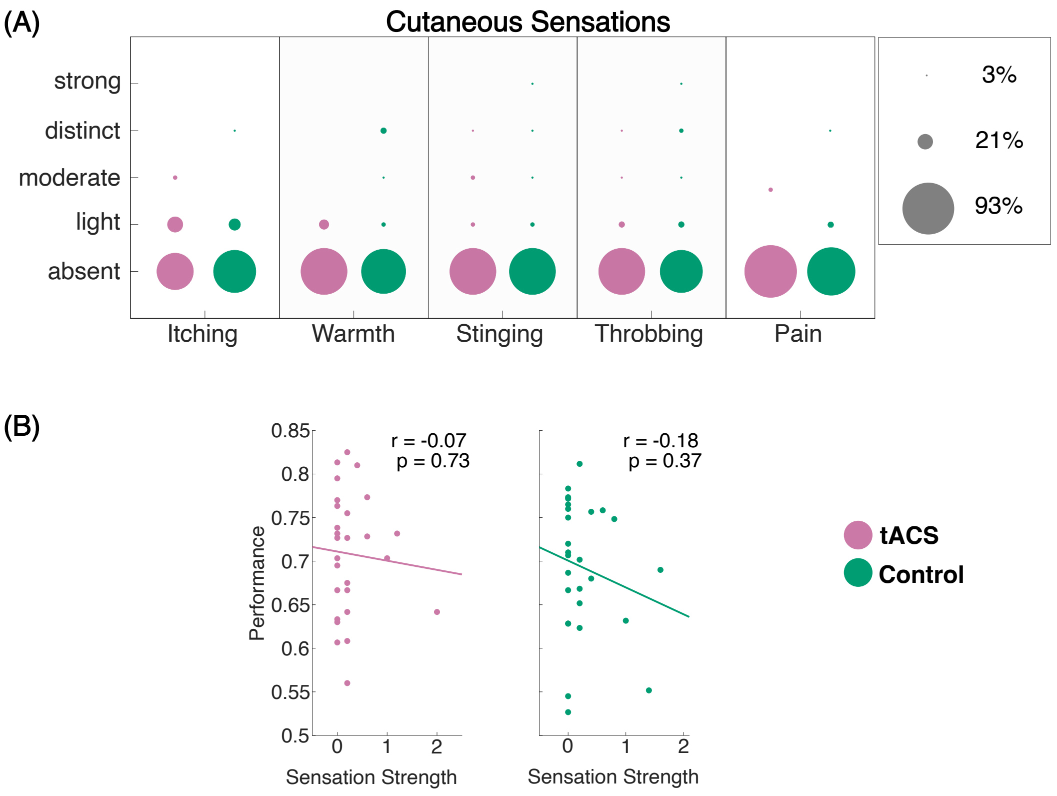


Figure 1: Unimodal experiment (red: tACS, green: Control). (A) Somatosensory artefacts of active tACS and active control condition, with no significant difference observed between conditions. Around 90% or more of the participant reported no cutaneous sensations. (B) Low correlation between cutaneous sensations and performance, further supporting the notion that tactile sensations did not significantly impact performance.

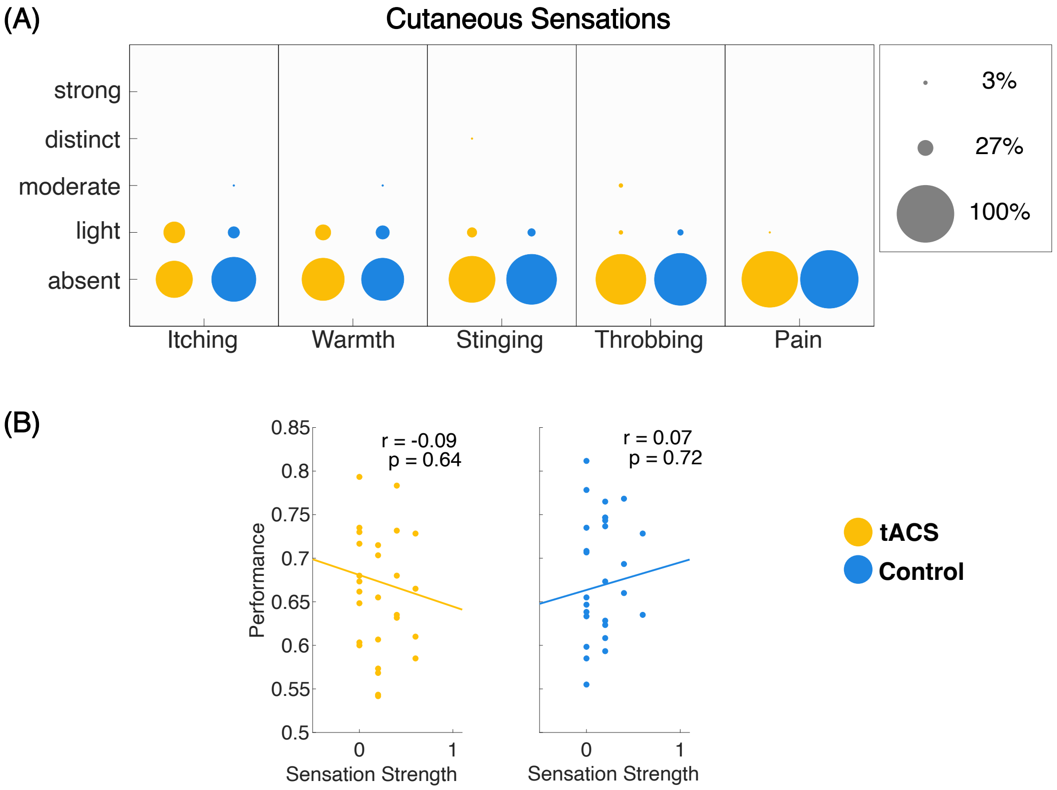


Figure 2: Crossmodal experiment (yellow: tACS, blue: Control). (A) Somatosensory artefacts of active stimulation and active control condition. No significant difference between conditions. Around 90% or more of the participant reported no cutaneous sensations. (B) Low correlation of cutaneous sensations and performance suggest no influence of tactile sensations on performance.
